# Validation of Illumina’s Isaac variant calling workflow

**DOI:** 10.1101/031021

**Authors:** Liudmila S. Mainzer, Brad A. Chapman, Oliver Hofmann, Gloria Rendon, Zachary D. Stephens, C. Victor Jongeneel

## Abstract

As the pace of implementing personalized medicine concepts increases, high-throughput variant calling on hundreds of individual genomes per day is a reality that will likely be faced by sequencing facilities across the country in the near future. While the scientific best practices for human variant calling workflows have been well defined, they also pose serious computational challenges at this high scale. Therefore, efforts in both academia and the private sector have focused on developing alternative workflows that may substantially reduce the computational cost per individual genome. Isaac is an “ultra-fast” variant calling workflow, designed by Illumina, Inc, and is claimed to be six times faster than BWA-GATK, with comparable sensitivity and specificity. This report is an independent review of Isaac, mainly focused on the accuracy of variant calls. We note that Isaac is indeed quite fast, and provide some benchmarks on a few hardware architectures. The overall conclusion from our analysis is that the Isaac workflow has undergone substantial improvement from version 01.14.11.27 to Isaac_2.0. The call accuracy is especially high on NA12878, however exomes tend to have a high fraction of false positive calls. We did not manage to reproduce the 99% sensitivity and specificity reported in the Illumina whitepaper, however that might be improved with further tweaking of the options. This report includes the information about some of the command-line parameters and documentation.

## Introduction

Genomic variant calling from raw high-throughput sequence data is a widely used procedure in both research and clinical settings. The current community standard (“BWA-GATK”) is a complex workflow that involves multiple steps, each requiring its own software tools and specific parameters. It can use a large amount of disk space for temporary files, and take a long time to compute. Therefore, there have been several efforts to develop better-performing sets of tools for variant calling, in terms of speed, accuracy, and ease of use. Isaac is an “ultra-fast” variant calling workflow, designed by Illumina, Inc. It is claimed to be six times faster than BWA-GATK, with comparable sensitivity and specificity (Isaac whitepaper, 2015). This report is an independent review of Isaac. Our main focus in this review is the accuracy of variant calls, which is the most important feature of any variant calling workflow. We note that Isaac is indeed quite fast, and provide some benchmarks on a few hardware architectures. However, measuring wall-time performance was not our priority, and we do not report any rigorous comparative analysis here.

Several groups have attempted to measure the discordance in variant calls among the many variant calling tools available to-date (Yi et al. 2014, Pabinger et al. 2014, Yu and Sun 2013, O’Rawe et al. 2013, Cornish and Guda 2014). Our focus here is not to provide a comparison between Isaac and other software packages. Instead, we focused on testing the accuracy of Isaac with datasets for which the “ground truth” for variant call is known, or at least agreed upon. These include synthetic data, data from the Genome in a Bottle (GIAB) consortium, and Illumina Platinum Genomes.

Illumina released several versions of Isaac in quick succession through 2015. Here we report our results, using the same data to test several different versions of the workflow.

The overall conclusion from our analysis is that the Isaac workflow has undergone substantial improvement from version 01.14.11.27 to Isaac_2.0. The call accuracy is especially high on NA12878, however exomes tend to have a high fraction of false positive calls. We did not manage to reproduce the 99% sensitivity and specificity reported in the Illumina whitepaper. That might be improved with further tweaking of the options. This report includes the information about some of the command-line parameters, and links to documentation.

## Caveats in the software design that affected our assessment

Isaac has several design caveats that impact the ability to evaluate and use it. We describe them briefly in the paragraphs below.

### Versions we evaluated

Isaac comes in two forms: (1) as commercially supported software that works directly off the output from an Illumina machine, and (2) a “developer” version. The former requires, apart from the usual FASTQ, the various Illumina files produced by the machine, and cannot be evaluated without them. Since our group does not have access to those, we focused on evaluating the “developer” version of Isaac. The latter is a two-step workflow that first performs alignment (versions 01.14.11.27, 01.15.04.01, Isaac_v2), followed by the variant calling step (version 1.0.7). Table 1 summarizes all experiments we have run. The software can be found in the following locations, as of November 25, 2015:

1. Isaac aligner 01.14.11.27 https://github.com/sequencing/isaac_aligner/tree/6b41f6d4985d069566a5c38d3d80c0d6ebb5841c
2. Isaac aligner 01.15.04.01 https://github.com/sequencing/isaac_aligner/commit/fd092ec20d82548a90c53eafdaed7d617328a6a4
3. Isaac_v2 https://github.com/Illumina/isaac2/
4. Isaac v2 workflow in Basespace: we used the app called "Isaac Whole Genome Sequencing v2 v2.0.0”
5. Variant caller https://github.com/sequencing/isaac_variant_caller

**Table 1.**
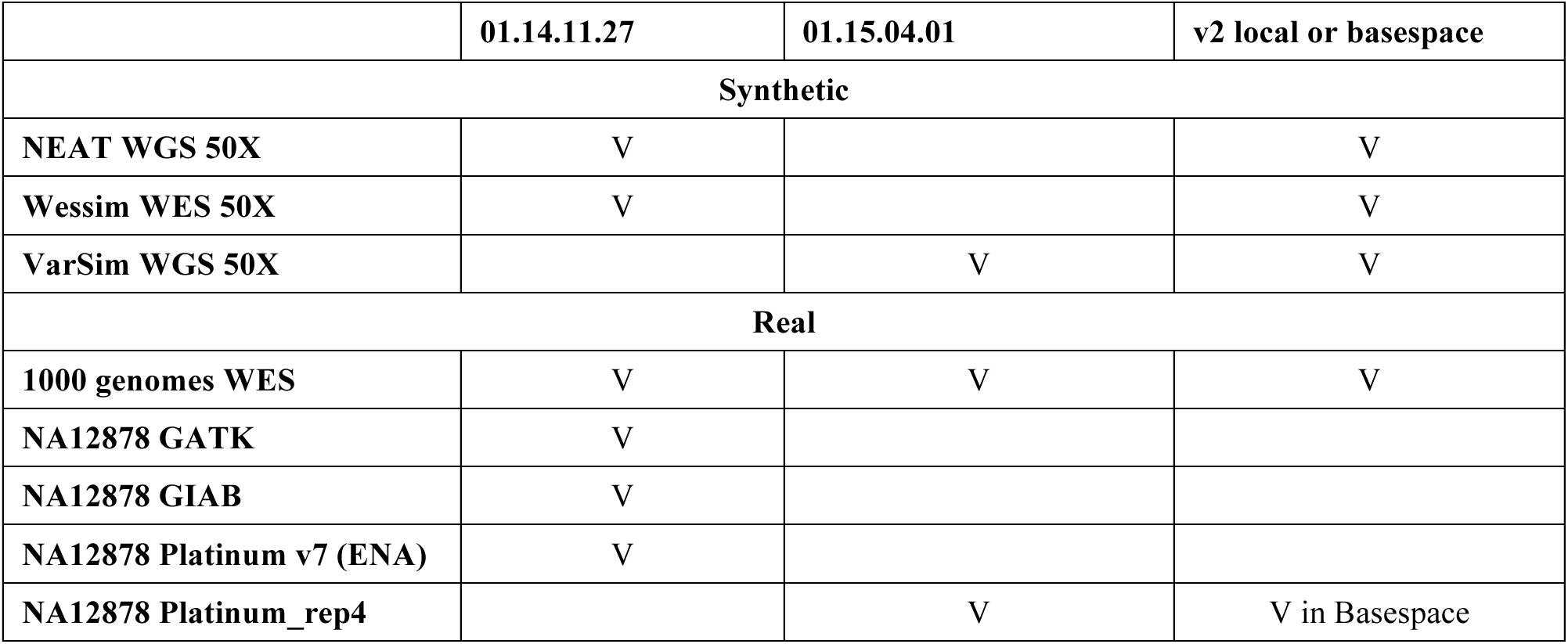
Summary of all experiments discussed in this report

### Input FASTQ names

Isaac expects the input FASTQ names to be in a certain format: `lane?_read?. fastq.` This probably allows it to grab the lane and read information directly from the name, for whatever purpose during the analysis. This could be accomplished by providing some user-set options on command line, but instead this requirement is hard-coded, and the tool simply will not run on FASTQ with any different kind of name. As a result, it cannot be used to perform high-throughput analysis on legacy data directly. An investigator must rename all files and perform his own book-keeping to keep track of which file was where.

### Input dbSNP

Isaac does not make use of a dbSNP, and there is no user option to specify one on command line.

### Read names

Isaac renames input reads; this is described in a manual on Github: https://github.com/sequencing/test/blob/master/markdown/manual.md. This means, a user cannot verify the Isaac alignment against a different tool, such as Novoalign (www.novocraft.com) or BWA (Li and Durbin 2009). Presumably, the reads are renamed with a sequential counter appended at the end of the new name. Thus, if a user resorts the output BAM by name, the reads should be in the same order as the original FASTQ, and correspondence can be established. However, if some reads were filtered out or collapsed as duplicates during an alignment, then the order is forever lost, and the original cannot be restored. One possible workaround could be to use Isaac output BAM files as input to BWA, and compare alignments that way. We did not perform that kind of analysis.

### Output folders

Isaac uses hardcoded folder structure. It will always create `Aligned/` and `Temp/` in the local folder wherefrom it was invoked, and will put all output into those two folders. In this situation, the user must either rename those folders by hand after the run, or `cd` into a pre-created output folder to invoke Isaac from there. Not a huge chore, but must be aware of it, as otherwise the output from different sequential runs will overwrite each other.

The output contents will be quite large: up to ~1TB for WGS human data (depth of coverage 50X) in `Temp/.` Final output, such as the alignment BAM and the associated statistics, is placed into `Aligned/.` Temporary run-time data are saved, obviously, into `Temp/,` which is not deleted after the run, so the user has a chance to inspect it.

The folder structure inside `Aligned/` seems to derive from some sequencing information (described in the GitHub manual): in our case the path to final BAMs was always `Projects/default/default.` Perhaps one can specify this information somehow as an input to Isaac, but that is not transparent.

### Documentation

There is no single comprehensive documentation for Isaac. Here are the sources we used.

1. For explanation for output VCF format and the variant filters, see the Isaac Whole Genome Sequencing v2 User Guide: http://support.illumina.com/content/dam/illumina-support/documents/documentation/software_documentation/basespace/Isaac-wgs-user-guide-15050954b.pdf
2. For example commands, output folder structure, aligner command line parameters see the GitHub manual: https://github.com/sequencing/test/blob/master/markdown/manual.md Please note that some of the options are not accurately described. For example, the `--variable-read-length` parameter takes values o`n|off`, not `=1|0,` as is implied in the manual. Experimenting with parameters is required.
3. For command line parameters in the variant caller, use the starling manual: invoke as`/Path/To/IsaacVariantCaller/libexec/starling2 -h` on command line. Please note that some options, such as `bsnp-ssd-no-mismatch, bsnp-ssd-one-mismatch` and `min-vexp` appear to be absent from starling command line documentation, as distributed by the Isaac variant caller package.
4. Many option settings are not known until the software is run, but can be gleaned from the vcf header.
5. Some options can be set in run.config.ini, but their names do not correspond exactly to those listed on GitHub manual or starling manual. Experimentation is required.
6. Some information in the HiSeq software Manual and user Guide could be helpful when trying to understand the parameter settings: https://support.illumina.com/content/dam/illumina-support/documents/documentation/software_documentation/has/hasuserguide/15041353b.pdf

## Methods

### The questions we asked

- Does Isaac find any variants?
- Does it find variants that should be there? If not, what is the rate of false negatives?
- If it does not find variants, can that be fixed with some parameter tuning?
- Does it find variants that should not be there? If so, what is the rate of false positives?

### Testbeds

The bulk of the work presented in this report was performed on the high-memory node of the Innovative Systems Lab (ISL2.0) at NCSA http://www.ncsa.illinois.edu/about/org/isl/ (Table 2). We chose this highly advanced machine, because we were running into difficulties running Isaac on any other, more conventional system (see the performance section).

**Table 2.** Computer systems we used to test Isaac for this report. Our tests were successful on the high memory machine administered by ISL2.0 at NCSA. However, tests resulted in errors and segmentation faults on two clusters: iForge (NCSA), and Biocluster (Institute for Genomic Biology http://help.igb.illinois.edu/Biocluster).

| Node | High Memory | iForge | Biocluster |
| --- | --- | --- | --- |
| System | Dell PowerEdge R920 | Intel Xeon Phi Development System in a cluster | SGI UV1000 Nodes in a cluster |
| CPU | 4 of Intel Xeon 2.6 GHz, yielding 48 dual-threaded cores, which allows for 96 threads | 2 x 6-core Intel Xeon x5670 "Westmere" 3.06 GHz | 384 of Intel Xeon 2.67 CPUs per node |
| RAM | 3 TB | 96 GB per node | 2 TB per node |
| Storage | 2 of 300 GB HDD, 6 of 1 TB HDD, 4 of 800 GB SSD, | No internal node storage, files saved over cluster's GPFS | No internal node storage, files saved over cluster's GPFS |
| Inter connect | 1 Gbps and 10 Gbps Ethernet | QDR Infiniband | 10 Gbps ethernet |

We also made a run on one node in BaseSpace (https://basespace.illumina.com/dashboard), but the hardware specifications are unknown. The BWA-GATK workflows were run on Blue Waters supercomputer (bluewaters.ncsa.illinois.edu). We also used AWS (https://bcbio-nextgen.readthedocs.org/en/latest/contents/cloud.html) to run BWA-GATK and Isaac: a single Amazon EC2 r3.8xlarge instance with 32 cores and 244Gb of memory, using an EBS provisioned SSD volume with 3000 IOPS.

### Runtime parameters and commands

All alignments were done against hg19, which was indexed by Isaac with seed length 32. All parameters for the alignment were default except specifying seed depth of the reference index, and the maximum RAM to use, which was specified as 2.999 out of the available 3 TB. All parameters for variant calling were default unless specified otherwise. All the 96 threads available on the machine were used. The box below lists the commands used during a typical test run.

~~~
<u>Alignment</u>
/PathTo_Isaac_install/bin/isaac-align
-r /Full/Path/sorted-reference.xml
-b /full/Path/FolderWithFASTQFiles
--base-calls-format fastq (or fastq-gz) -m 2999 --seed-length 32
-j 96
It was sometimes necessary to use:
-variable-read-length on
<u>Variant Calling</u>
mkdir VarCallFolder
cd VarCallFolder
cp /PathTo_Isaac_VariantCallerFolder/etc/ivc_config_default.ini ./config.ini
/PathTo_Isaac_VariantCallerFolder/bin/configureWorkflow.pl
--BAM=/PathToAlignedBAM/input.BAM --ref=/PathToReferenceFile/hg19.fa
--config=./config.ini --output-dir=./myAnalysis
cd ./myAnalysis
make -j 96
~~~

When needed, we ran the GATK best-practices workflow using the following commands and parameters:

~~~
<u>Alignment</u>:
bwa mem -k 32 -t 30 -I 300,30 -R ‘read-group-string’ /Path/To/Reference/Index /Path/To/leftreads /Path/rightreads | samblaster -o /Path/To/Output/SAM
or
Novoalign -g 40 -x 5 -I PE 175,50 -r Random -hdrhd off -v 120
~~~

~~~
<u>Convert to BAM:</u>
samBAMba view -t 32 -f BAM -S /Path/To/Output/SAM -o /Path/To/Output/BAM
<u>Sort BAM:</u>
novosort --threads 32 --index /Path/To/Output/BAM -o /Path/To/Output/BAM.sorted
<u>Gatk Create realignment targets</u>
-T RealignerTargetCreator
-R genome.fa -known dbsnp_135.hg19.vcf.gz
-I BAM.sorted -o BAM.sorted.realign-list
~~~

~~~
<u>Realignment:</u>
-T IndelRealigner
-R genome.fa -known dbsnp_135.hg19.vcf.gz
-targetIntervals BAM.sorted.realign-list
-I BAM.sorted -o BAM.sorted.realigned
<u>Base score recalibration:</u>
-T BaseRecalibrator
-R genome.fa --knownSites dbsnp_135.hg19.vcf.gz
-I BAM.sorted.realigned.BAM --out BAM.sorted.realigned.recal-report.grp
-nct 32
<u>Print reads:</u>
-T PrintReads
-R genome.fa
-BQSR BAM.sorted.realigned.recal-report.grp
-I BAM.sorted.realigned.BAM --out BAM.sorted.realigned.recalibrated
-nct 32
<u>Variant calling:</u>
-T UnifiedGenotyper
-R genome.fa
-I BAM.sorted.realigned.recalibrated -o raw.vcf
-glm BOTH
--output_mode EMIT_VARIANTS_ONLY
-A Coverage
-A AlleleBalance
-dcov 250
-rf BadCigar
-nt 8 -nct 4
~~~

## Concordance measurement

To measure the concordance of variant calls, we would have liked to perform comparisons of vcf files as well as the alignment BAMs, as the latter can sometimes help explain discordance. Since Isaac changes the read names, the BAM comparison is therefore impossible. Instead, we spot-checked the Isaac aligned BAMs in a genome browser.

Additionally, Isaac outputs variants in a GVCF, which conforms to the VCF4.1 specifications but also includes non-variant sites. This can cause conventional VCF comparison tools to report incorrect results. Thus, we extracted the variants from the Isaac’s output gvcf. The Isaac documentation recommends using `extract_variants` from `gvcf-tools.` After using it, we found that it performs the same function as a linux `awk` command, but slower. Indeed, one has to use awk afterwards anyway to inspect the variants that did not pass the various filters. Thus, we used the following commands to go from Isaac’s GVCF to a VCF that can be used in comparison exercises:

~~~
awk ‘$1!~/^^^#/ && $5!~/\./ {print $0}’ sorted.genome.vcf > sorted.genome.vcf.ALT
awk ‘$7~/PASS/ {print $0}’ sorted.genome.vcf.ALT > sorted.genome.vcf.ALT.PASS
~~~

The first command produces the list of all variants, whether or not they passed the filters (discussed in the Results section). The second command produces the list of only those variants that have passed the filters. Afterwards, we manually inspected the vcfs and used existing tools, such as `vcf-compare`, as well as our own `perl` and `python` scripts. The “confident regions” from the Illumina Platinum Genomes v8 were also used in the analysis. The minimum length of a targeted region (in WES) or a confident region (from the Platinum set) that was considered in the concordance analysis was 20 nt.

## Generating synthetic WGS with NEAT

To test Isaac on synthetic whole human genome with known variants (WGS), we produced one synthetic dataset using NEAT (https://github.com/zstephens/genReads1) at 50X depth. The variants were inserted at random, at the average rate of 0.00034. The simulated fragment length was 300 nt, standard deviation of 30 nt, read length of 100 nt. The sequencing error rates were modeled after the data generated at the local sequencing facility (http://www.biotech.uiuc.edu/htdna), and were inserted at the rate of 0.1%. The software produces the synthetic reads, a “golden” vcf containing the variants synthetically inserted into the reference, and the “golden” sam containing the “true” read alignment based on the reference loci where the reads were generated from.

## Generating synthetic WES with Wessim

To test Isaac on synthetic whole human exome with known variants (WES), we produced one synthetic dataset using Wessim (Kim et al. 2013):

1. We ran a standard GATK workflow on ERR250949 human exome from the 1000 genomes project, detected variants and segregated out those belonging to chromosome 1.
2. We generated some random variants as well, using Genome Smasher (10% insertions, 5% deletions and the rest are SNPs; regions with repetitive Ns were avoided).
3. Then we combined those two sets of variants together and inserted them into hg19 reference using GATK FastaAlternateReferenceMaker. Only variants located within exonic regions identified according to hg19 annotation, were considered in concordance measurement.
4. Finally, we used that mutated reference to simulate whole exome sequencing on chromosome 1 using Wessim, generating 100-nucleotide paired-ended reads at 50X depth.

## Results

### Synthetic WGS 50X, generated with NEAT

#### Isaac 01.14.11.27 detects no variants

It was desirable to test Isaac’s variant detection against a known “ground truth”, in order to evaluate its accuracy unequivocally, and independently of the properties of other variant callers. Thus, we generated a synthetic WGS dataset based on hg19 at 50X using NEAT, and compared Isaac’s results to the list of variants that were inserted by the read simulator. Unfortunately, Isaac 01.14.11.27 did not detect any variants at all. The final GVCF contained no entries in the ALT column, only dots.

We hypothesized that perhaps our variants were filtered out by the variant caller, so we relaxed the following filters that have to do with local read depth and read mapping quality (by the way, these parameters have different names in the starling manual and Isaac variant caller configuration file):

~~~
isSkipDepthFilters=1
maxInputDepth=-1
depthFilterMultiple=1000
minMapq=0
minGQX=0
~~~

This had no effect. Inspecting the BAM in genome browser indicates that some expected variants do in fact get detected in alignment – but are not reported by the variant caller. For example, this variant was inserted into the dataset when simulated reads were created, and is present in the “golden” vcf (line abbreviated):

~~~
chr1        240578        .        G        A        .
~~~

However, Isaac variant caller reports nothing for the respective region (lines abbreviated):

~~~
#CHROM    POS    ID          REF     ALT      QUAL    FILTER
chr1             240547   .             T          .              .            HighDPFRatio
chr1             240549   .             A          .              .            HighDPFRatio
chr1             240553   .             G          .              .            HighDPFRatio
chr1             240583   .             A          .              .            HighDPFRatio
chr1             240596   .             C          .              .            HighDPFRatio
~~~

Meanwhile, the same variant is clearly visible in Isaac’s alignment (Figure 1).

**Figure 1.**
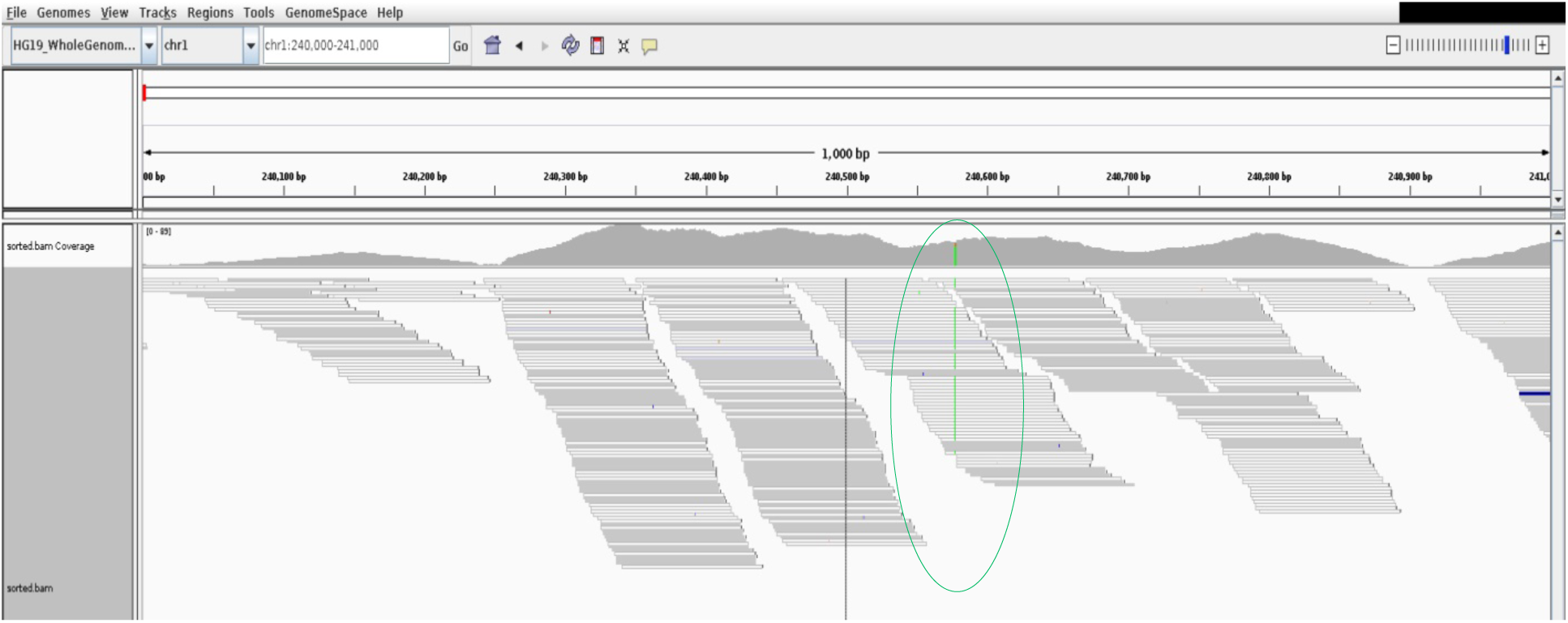
Isaac’s alignment on NEAT synthetic data; snapshot from the genome browser IGV. The variant at Position chrl:240578 is highlighted in green.

Both Novoalign-GATK, and BWA-GATK work fine on this kind of data and report excellent concordance of variants (Figure 2).

**Figure 2.**
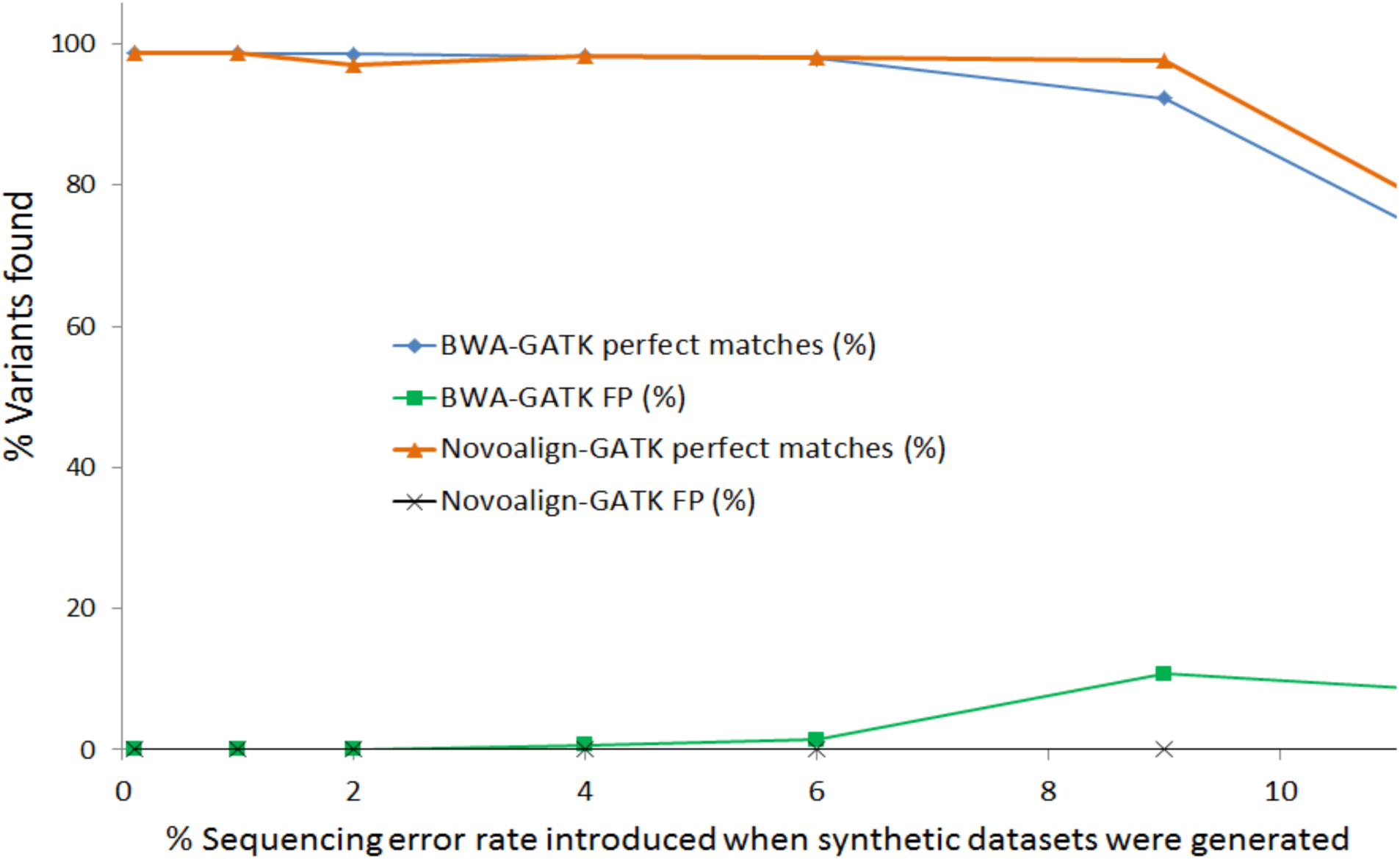
Results of running the standard GATK workflows on the NEAT synthetic WGS, 50X. Reads were simulated with varying sequencing error rates (abscissa). Perfect matches denote the number of variants found by the GATK workflow that corresponded to the “golden” vcf. False positives (FP) denote the number of variants found by the GATK workflow, but which were not inserted into the data at the time of simulation.

#### Isaac v2 provides concordance of up to 97% on NEAT data

Release of Isaac v2 aligner resulted in significant improvement. While the earlier version detected no variants at all, v2 provides fairly high concordance (Table 3).

**Table 3.**
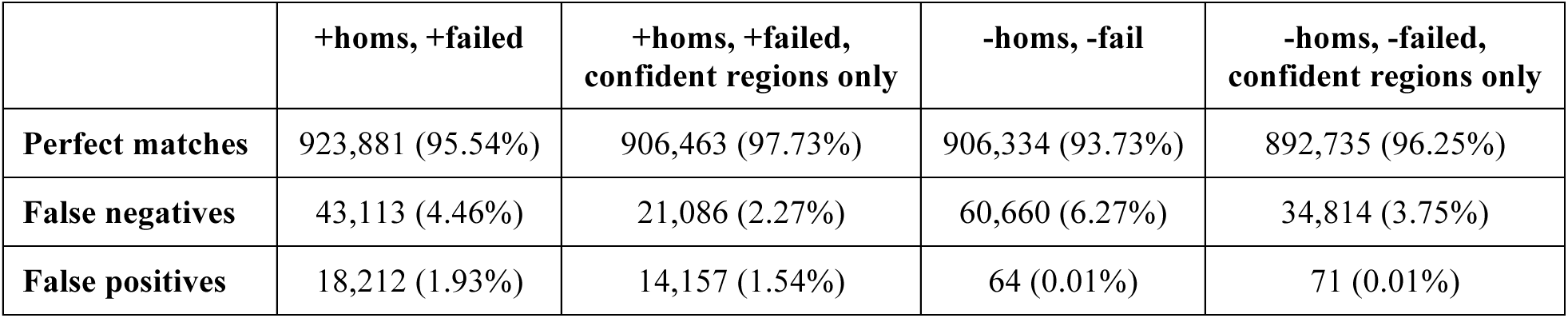
Concordance between variants called using Isaac_v2 and the variants inserted into the synthetic reads by NEAT. Measurements were done both ways: on all reported variants, including those that are homozygous and failed filtration (+homs, +failed), and also on the subset excluding those two categories (-homs, -failed).

|  | +homs, +failed | +homs, +failed,<br>confident regions only | -homs, -fail | -homs, -failed,<br>confident regions only |
| --- | --- | --- | --- | --- |
| **Perfect matches** | 923,881 (95.54%) | 906,463 (97.73%) | 906,334 (93.73%) | 892,735 (96.25%) |
| **False negatives** | 43,113 (4.46%) | 21,086 (2.27%) | 60,660 (6.27%) | 34,814 (3.75%) |
| **False positives** | 18,212 (1.93%) | 14,157 (1.54%) | 64 (0.01%) | 71 (0.01%) |

### Synthetic WGS 50X, generated with VarSim

We tested Isaac_v2 on synthetic data generated with VarSim, as an alternative to our NEAT synthetic reads. The VarSim data were downloaded from the developer’s repository. The Isaac alignment aborted with the message:

~~~
Opened gz fastq stream on lane1_read1.fastq.gz
Opened gz fastq stream on lane1_read2.fastq.gz
ERROR: Thread: 1 caught an exception first:Invalid
argument:Isaac_v2/isaac2/src/c++/include/io/fastqReader.hh(170): Throw in function
InsertIt isaac::io::fastqReader::extractBcl(const Isaac::flowcell::ReadMetadata&,
InsertIt) const [with InsertIt =___gnu_cxx::___normal_iterator<char*, std::vector<char>
>]
Dynamic exception type:
boost::exception_detail::clone_impl<isaac::io::FASTQFormatException>
std::exception::what: Invalid oligo M found in lane1_read2.fastq.gz at offset
5966430356
:Invalid oligo M found in lane1_read2.fastq.gz at offset 5966430356
~~~

Looking back at the synthetic sequences, they did indeed contain the “nucleotide” M:

~~~
grep -n M VarSim/lane1_read2.fastq
94111002:
CAGCTTGCTCTTCATTAGCGCTACATAGCTGMCTTATTATTCGTGGTCCCGTATGACCCCCTGATCATTTTCCCTGAGGGTGCATA
TTTATTCACTAACT
103350290:
TGTTTTTCATTTTCTTGATTTATTTCTGAATTCAGCTTGCTCTTCATTAGCCCTACATAGCTGMCTTATTATTCGTGGTCCCCTAT
GACCCCCTGGTCAT
116458522:
CATTAGCGCTACATAGCTGMCTTATTATTCGTGGTCCCCTATGACCCCCTGATCATTTTCCCTGAGGGTGCATATTTATTCACTAA
CTATGTTACAATCA
123122630:
AATTCAGCTTGCTCTTCATTAGCGCTACATAGCTGMCTTATTATTCGTGGTCCCCTATGACCCCCTGATCATTTTCCCTGAGGGTG
CATATTTATTCACT
128669782:
CTTGATTTATTTCTGAATTCAGCTTGCTCTTCATTAGCGCTACATAGCTGMCTTATTATTCGTGGTCCCCTATGACCCCCTGATCA
TTTTCCCTGAGGGT
~~~

### Wessim synthetic WES, 50X

#### Isaac 01.14.11.27 generates a high number of false positives

We produced one synthetic dataset using Wessim (Kim et al. 2013), which is known to closely simulate the properties of Illumina exome sequencing data. The Isaac version 01.14.11.27 did report some variants on this dataset, but with a very high rate of false positives (at least 251%) and false negatives (at least 28%). Out of 7,314 artificially inserted variants (all within exonic regions, according to hg19 annotation; see salmon ellipse on Figure 3), 2,048 were undetected (false negatives). Out of 23,638 variants that Isaac claimed to be present in this dataset (cyan ellipse on Figure 3), 18,374 were not a part of the synthetic set (false positives).

**Figure 3.**
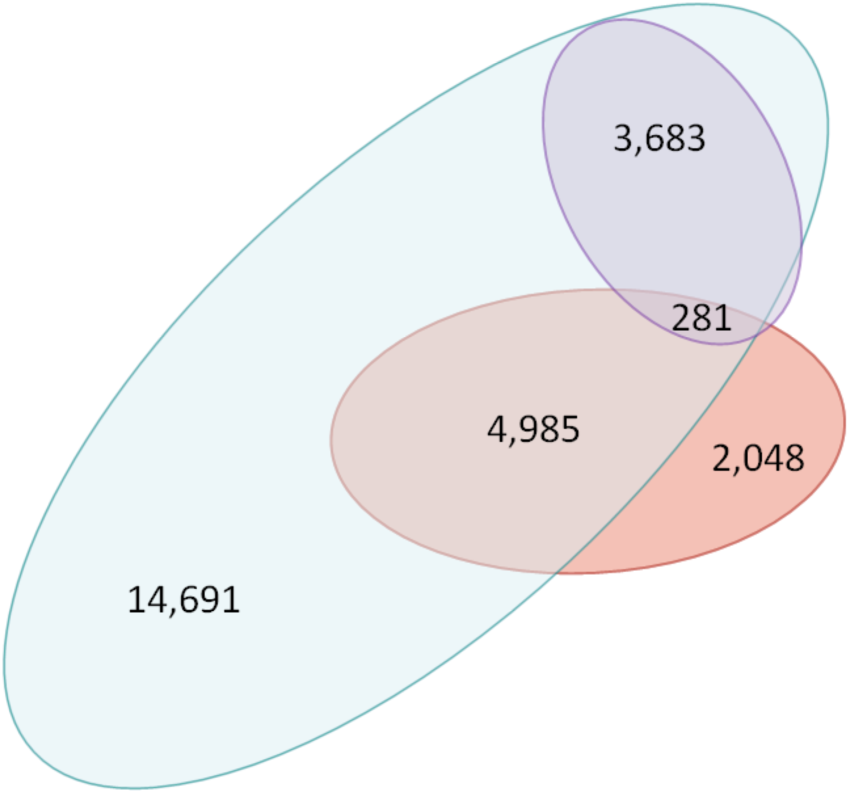
Concordance statistics on the synthetic WES chr1 data generated by Wessim. Out of 7,314 simulated variants, 5,266 were detected by Isaac, but only 281 of those actually passed the default filters. Salmon ellipse: synthetic variants. Cyan ellipse: variants detected by Isaac. Purple ellipse: variants that were detected by Isaac and passed the default filters.

Looking at Isaac’s output vcf more closely, it appears that few of the claimed variants pass the default filters (purple ellipse on Figure 3). Those filters are listed in Table 4, copy-pasted from Isaac’s vcf header.

**Table 4.**
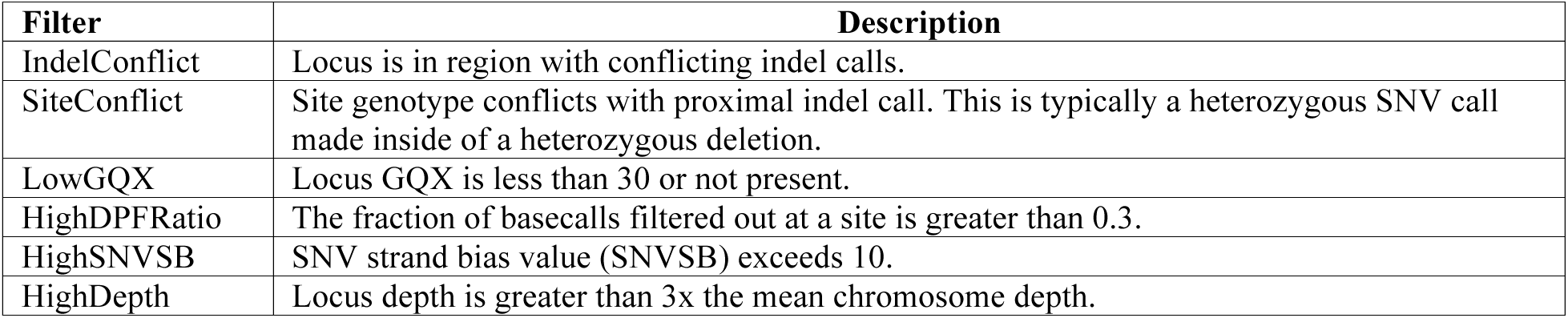
Default filters used by Isaac, copy-pasted from an output vcf. In the particular case of Wessim-generated reads, most of the inserted variants fail to pass the LowGQX or HighDepth filters (Table 4). GQX has the meaning of genotype quality score, generalized to both variant and non-variant loci. An example of such filtered variants is displayed on Figure 4.

**Table 5.** Synthetic variants that were found by Isaac but did not pass the default filters, broken down by the filter.

| Filter | Number of variants |
| --- | --- |
| HighDepth | 4591 |
| HighDepth, HighSNVSB | 2 |
| HighDPFRatio | 12 |
| HighDPFRatio, HighDepth | 80 |
| LowGQX | 217 |
| LowGQX, HighDepth | 1 |
| LowGQX, HighDPFRatio | 21 |
| LowGQX, HighDPFRatio, HighDepth | 16 |

Filtering variants based on the sequencing depth and quality score is, of course, normal, and expected. The problem lies not with the fact that variants get filtered out, but with the extremely small number of passed variants that we know have actually been inserted into the data during the process of read simulation.

The cause of this is the way Wessim simulates exome sequencing: reads are grabbed from the reference, in and around target regions. There are usually reads present that do not strictly belong to exons, and they will usually have coverage issues. Once we focus only on the 3,792 synthetic variants that belong strictly to the targeted regions (defined by the Agilent SureSelect bed file, and only considering regions with length > 100), actually Isaac detects 98.55% of them (3,737 variants), but only 15 of them pass the filters. Additionally, Isaac finds 2,781 extra variants that should not be there (73.34%, only 36 passed filters; false positives) within the targeted regions and does not find 55 (1.45%; false negatives) variants that should in fact be present within the dataset. In other words, Isaac does find strictly exonic variants within the targeted regions, but its filters are a bit stringent for this dataset, and need some tuning to work properly. This tuning is difficult: when the filters are relaxed, the true variants will pass filtration, but the false positive variants will pass it too.

Finally, Isaac assigned 993 variants to chromosomes other than chr1 (all regions), even though the input reads were simulated for chr1 only. This is normal, due to some sequence similarities between chromosomes. None of these variants passed the default filters.

Based on these data it appears that Isaac 01.14.11.27 can be quite sensitive to variants, but also has an incredibly high rate of false positives, which is only somewhat compensated by stringent variant filters.

**Figure 4.**
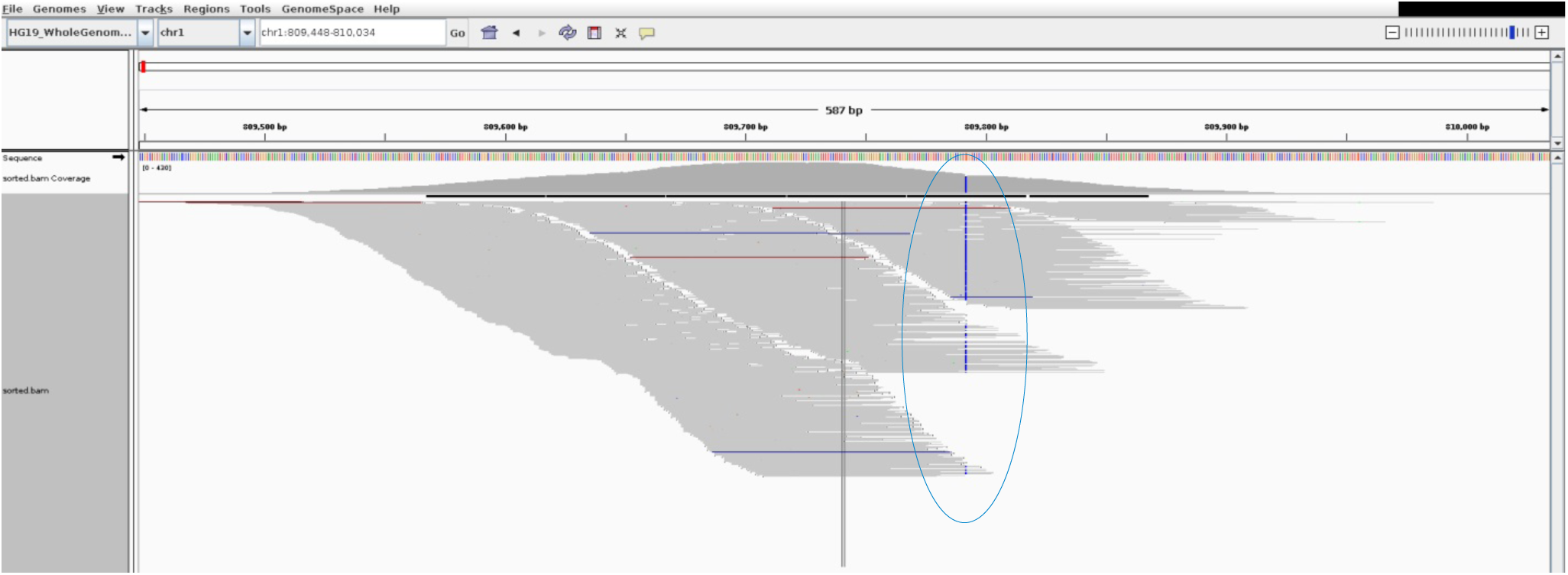
Example of two synthetic variants that were detected but did not pass the filters. This is a screenshot of IGV genome browser zoomed in on loci `chr1: 809, 792` and `chr1: 809, 956` within an output BAM from Isaac aligner. The first synthetic variant (G→C) is highlighted with blue in the figure, but is filtered out as HighDepth. The second variant (T*→*A) is on the tail edge of the simulated exon and is only covered by two reads, so it is barely visible, but is nonetheless present in Isaac vcf and marked as LowGQX.

#### Isaac v2 does not eliminate false positives on Wessim data

Since Isaac v2 aligner improved the results on synthetic NEAT data, we applied it to Wessim data as well. Unfortunately, the number of false positives was still extremely high (Table 6).

**Table 6.**
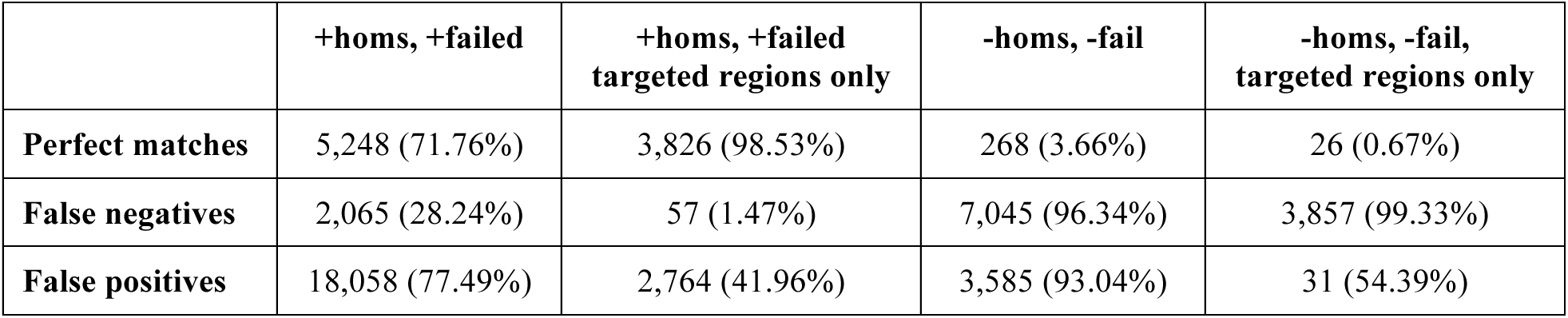
Concordance between variants called using Isaac_v2 and the variants inserted into the synthetic reads by Wessim. Measurements were done both ways: on all reported variants, including those that are homozygous and failed filtration (+homs, +failed), and also on the subset excluding those two categories (-homs, - failed). Concordance within the Platinum Confidence regions of hg19 is even worse than within targeted regions (73.85% perfect matches, 26.15% false negative, 77.66% false positives on +homs, +failed).

|  | +homs, +failed | +homs, +failed<br>targeted regions only | -homs, -fail | -homs, -fail,<br>targeted regions only |
| --- | --- | --- | --- | --- |
| **Perfect matches** | 5,248 (71.76%) | 3,826 (98.53%) | 268 (3.66%) | 26 (0.67%) |
| **False negatives** | 2,065 (28.24%) | 57 (1.47%) | 7,045 (96.34%) | 3,857 (99.33%) |
| **False positives** | 18,058 (77.49%) | 2,764 (41.96%) | 3,585 (93.04%) | 31 (54.39%) |

### 1000 human genomes, WES data

#### Isaac 01.14.11.27 generates high number of false positives

To eliminate any uncertainty in our evaluation that may have resulted from incorrect read simulation by Wessim, we proceeded to test Isaac on real datasets. We used whole exome sequencing dataset ERR250440 from the 1000 genomes project. The reads were aligned against hg 19 using Novoalign and then run through the best practices GATK workflow.

The resultant vcf was used as a standard to compare Isaac’s vcf against. Once again, some of the same variants were detected, but most were marked as not passing the filters, mainly due to high coverage (Figure 5).

**Figure 5.**
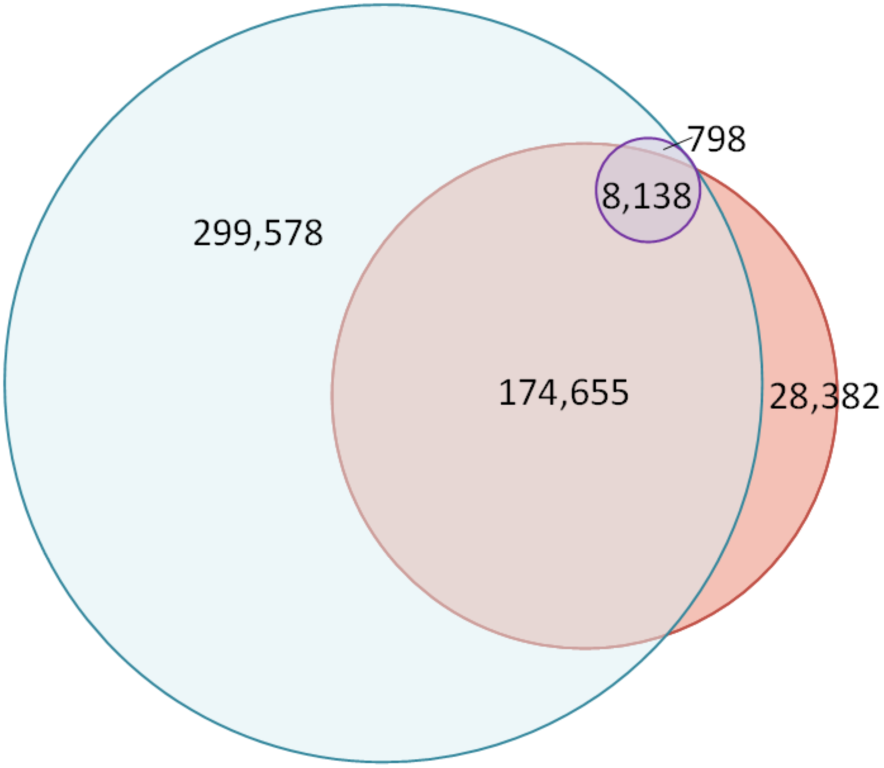
Concordance statistics on the variants detected with novoalign-GATK and those detected by Isaac. Out of 211,175 variants detected by GATK (salmon circle), 182,793 (86.56%) were also found by Isaac, but only 8,936 of those passed the filters. A large number of false positives is observed again (300,376), but only 798 of them pass the filters. Cyan circle: variants found by Isaac. Purple circle: variants found by Isaac that also passed filtration.

The number of false negative variants is still high (13.44%), and interestingly some of these are clearly visible in alignment, but not reported by Isaac. For example, the box below lists neighboring locations in Isaac’s and GATK’s vcfs, and matches the region displayed in the genome browser screenshot on Figure 6. All six variants listed in the box appear present in the alignment: their respective loci are highlighted in color within the box.

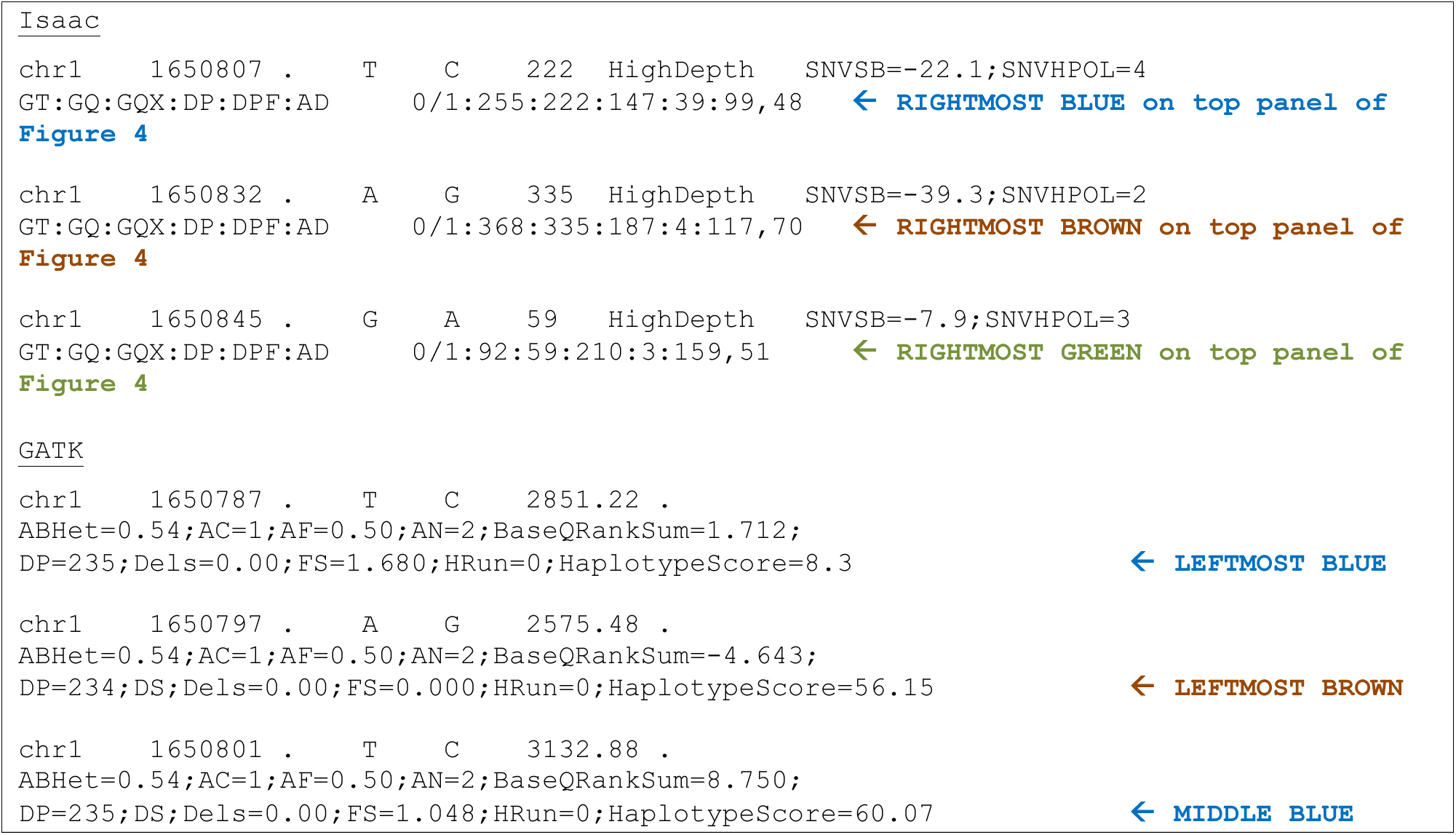

**Figure 6.**
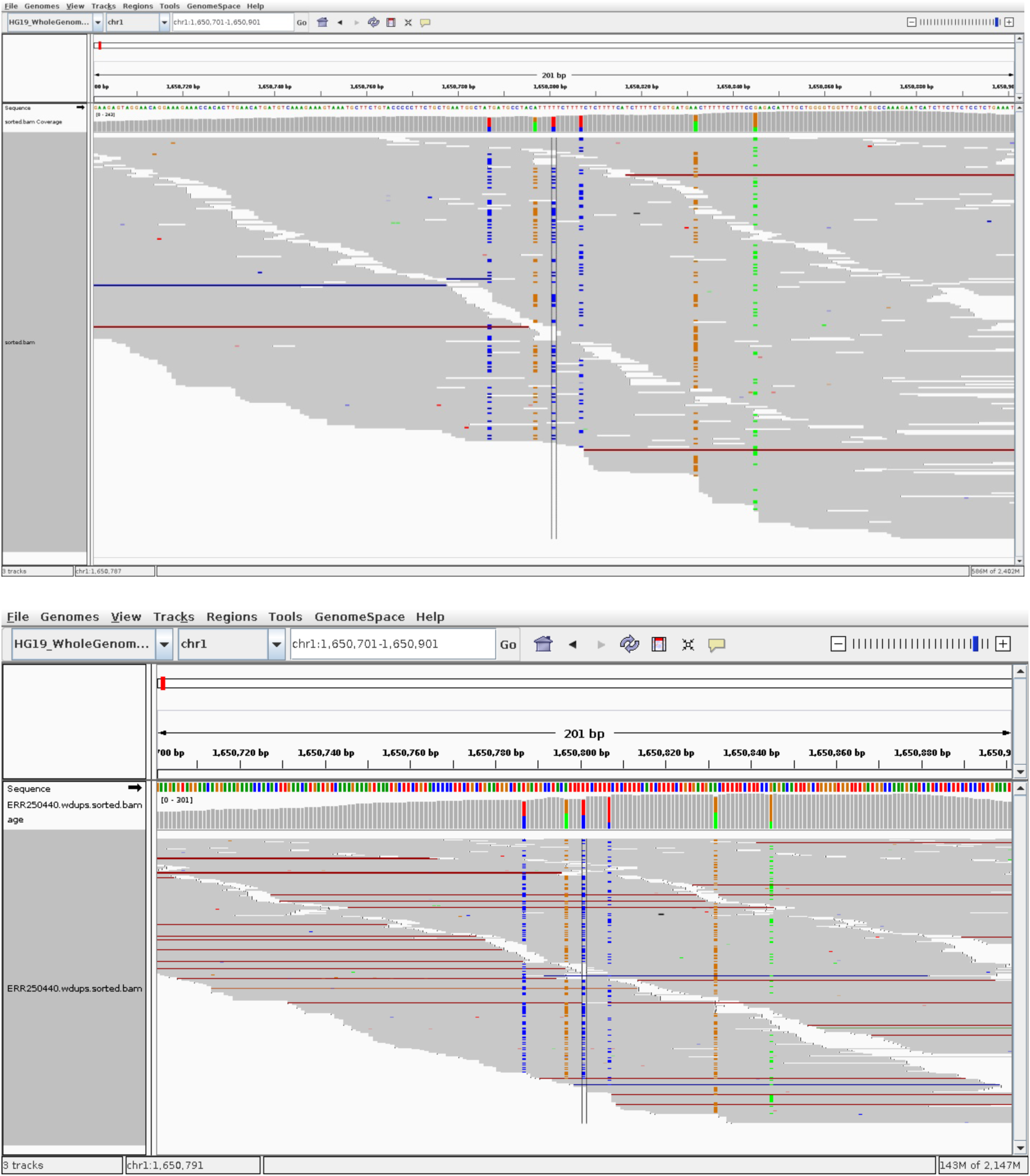
Example of variant mismatch between GATK and Isaac. Top panel is a screenshot of IGV genome browser zoomed in on the region `chr1:1, 650, 701–1, 650, 901` within an output BAM from Isaac aligner. Bottom panel is the same location within an output BAM from Novoalign. .

At first it might look like reads are misaligned, and single variants became split into two, and only one half got called. For example, variant `chr1:1650807` got called by Isaac, but `chr1:1650801` and `chr1:1650787` did not, even though they seem to have similar characteristics. However, the alignment performed by Novoalign looks extremely similar in the same region (Figure 6, bottom panel). Perhaps the variant caller is the culprit?

#### Isaac 01.15.04.01 and Isaac v2 both report high number of false positives on WES

Newer versions of Isaac do not eliminate the high false positives rate, although the number of perfect matches within the Platinum v8 confident regions is quite high (Table 7).

**Table 7.** Concordance between variants called using Isaac 01.15.04.01 and Isaac_v2 (second % number, where applicable), and the variants detected by Novoalign-GATK in the WES reads from the ERR250440 sample (1000 genomes project). Measurements were done both ways: on all reported variants, including those that are homozygous and failed filtration (+homs, +failed), and also on the subset excluding those two categories (-homs, -failed)

|  | +homs, +failed | +homs, +failed<br>Illumina targeted<br>regions only | +homs, +fail,<br>Platinum v8 confident<br>regions only | -homs, -fail |
| --- | --- | --- | --- | --- |
| **Perfect matches** | 184,511 (87.42%, 87.52%) | 34,336 (90.56%) | 172,467 (95.66%, 95.97%) | 8,430 (3.99%) |
| **False negatives** | 26,556 (12.58%, 12.48%) | 3,579 (9.44%) | 7,820 (4.34%, 4.03%) | 202,637 (96.01%) |
| **False positives** | 306,873 (62.45%, 66.37) | 11,064 (24.37%) | 290,573 (62.75%, 66.41%) | 863 (9.29%) |

### NA12878: paired-ended data from GATK and GIAB, and mate-pair data from ENA

Finally, to test variant detection for whole genome sequencing, we used three datasets derived from the Illumina Platinum genome NA12878. One dataset was sequenced by the Broad Institute http://gatkforums.broadinstitute.org/discussion/1292/which-datasets-should-i-use-for-reviewing-or-benchmarking-purposes to produce high-coverage (>60X) paired ended reads hosted by the NCBI ftp://ftp-trace.ncbi.nih.gov/1000genomes/ftp/technical/working/20101201_cg_NA12878/. The second set was downloaded from Genome In A Bottle (GIAB) consortium. The third was sequenced using long insert mate pair library (2000 nt fragment length) and hosted by the ENA http://www.ebi.ac.uk/ena/data/view/ERP002490.

The results are definitely better than for any dataset above. Was Isaac tuned to performed particularly well on NA12878? The number of false positives and false negatives is still noticeable in some cases, especially the mate pair ENA dataset (Table 8). This might be improved by setting the fragment length correctly, but it is not transparent from the documentation how to do that.

For comparison, Table 8 also includes the concordance stats on the GIAB data analyzed using BWA-GATK HaplotypeCaller.

**Table 8.**
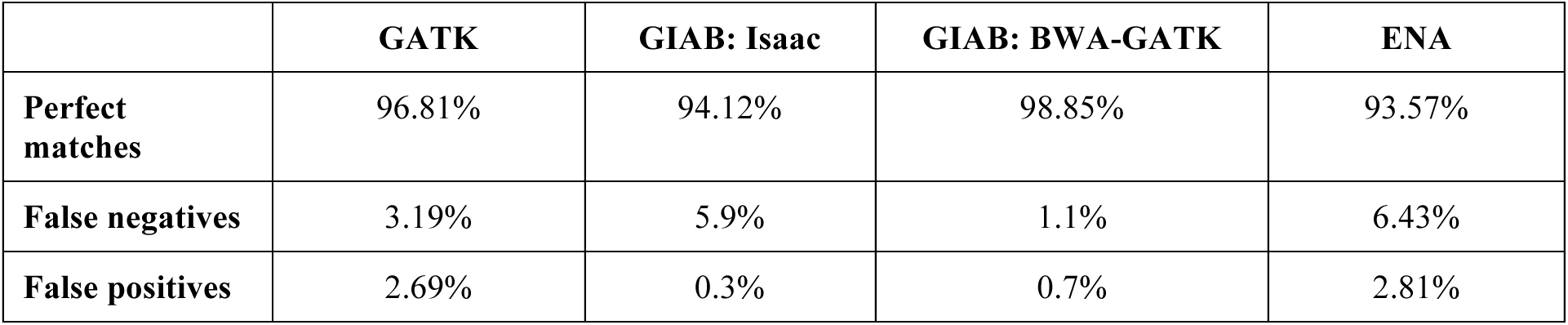
Concordance for variants called using Isaac 01.14.11.27 and BWA-GATK HaplotypeCaller on NA12878 datasets of various origins. Measurements were made on all reported variants, including those that are homozygous and failed, but only inside the confident regions. Variants called on the datasets sourced from GATK and ENA were compared against the Platinum set v7. The variants called on datasets sourced from GIAB were compared to variants provided by GIAB.

|  | **GATK** | **GIAB: Isaac** | **GIAB: BWA-GATK** | **ENA** |
| --- | --- | --- | --- | --- |
| **Perfect matches** | 96.81% | 94.12% | 98.85% | 93.57% |
| **False negatives** | 3.19% | 5.9% | 1.1% | 6.43% |
| **False positives** | 2.69% | 0.3% | 0.7% | 2.81% |

### NA12878_rep4 recommended by Illumina team

To test for the best concordance, the Illumina team shared with us in BaseSpace a dataset comprised of NA12878 sequenced on two lanes, paired-ended reads 126 nt long, 40X WGS. They called it NA12878_rep4 and recommended we compare the results against Illumina Platinum VCFs version 8.

We downloaded these data to the high-memory machine and ran the analysis locally using Isaac 01.15.04.01, with results comparable to those reported by Isaac 01.14.11.27 on the high coverage data from GATK. Isaac v2 was run on the NA12878_rep4 in BaseSpace, with very similar results (Table 9).

**Table 9.** Concordance between variants called on NA12878_rep4 using Isaac 01.15.04.01or Isaac v2, and the Platinum v8. Measurements were made on all reported variants, including those that are homozygous and failed filtration (+homs, +failed).

|  | **Isaac 01.15.04.01, local run** |  | **Isaac v2, in BaseSpace** |  |
| --- | --- | --- | --- | --- |
|  | **+homs, +failed** | **+homs, +failed**<br>**confident regions only** | **+homs, +fail** | **+homs, +fail,**<br>**confident regions only** |
| **Perfect matches** | 4,088,290 (91.68%) | 3,886,538 (96.15%) | 4,099,301 (91.93%) | 3,895,220 (96.36%) |
| **False negatives** | 370,778 (8.32%) | 155,675 (3.85%) | 359,767 (8.07%) | 146,993 (3.64%) |
| **False positives** | 346,085 (7.80%) | 91,669 (2.30%) | 35,020 (7.95%) | 94,363 (2.37%) |

## Performance

### Installation

Isaac installs fairly easily on a linux box with standard, modern OS distribution and libraries. It does require `boost` and `gnuplot` as prerequisites.

### Performance benchmarks

It takes a long time to index the reference: 8-12 hours depending on the number of available cores. This only needs to be done once.

On the Dell machine with 48 dual-threaded cores and 3 TB of RAM, the alignment takes ~80-120 minutes (depending on the version and OS activity), and variant calling in ~30 minutes on human WGS, 50X coverage. Performance definitely was degraded when sharing the machine with other bioinformatics applications, and sometimes Isaac crashed in those cases. On less powerful servers the alignment can take up to 4.5 hours and variant calling ~ 40 minutes. In BaseSpace, the entire workflow took 13.5 hours on a single server. In AWS, Isaac alignment took 4.5 hours, and variant calling took 45 minutes.

Isaac aligner seems to use all available computational resources on the host machine during the run (Figure 7). According to Isaac whitepaper, the alignment consists of three distinct phases, which manifest in performance profiles as well. The first phase generates mapping position candidates via seed-based search, and finds exact matches on a stream of input data, which seem to be kept in memory. The second phase finds best mapping among all selected candidate mappings and determines alignment scores. This phase is highly i/o intensive and CPU intensive. Finally, duplicates are found and removed - a process that seems to happen in RAM as much as possible. The aligner logs also mention realignment around gaps during this phase.

**Figure 7.**
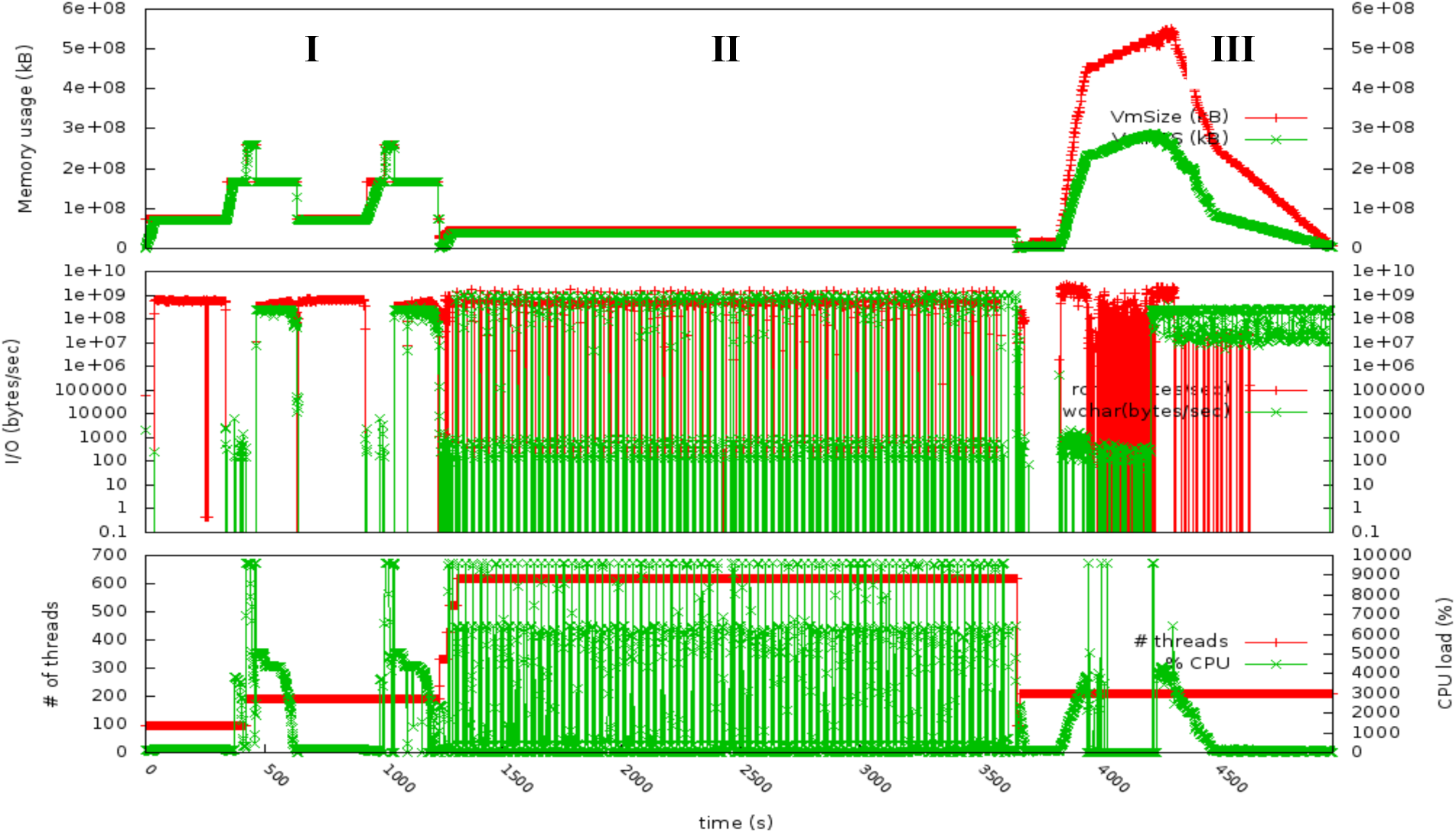
Performance profile on Isaac aligner, made by parsing /proc/PID every second. Top panel: RAM utilization measurements, specifically Vm size (red) and resident set size (green). The roman numerals mark the three computational phases discussed in the text. Middle panel: the rate of data reads (red) and writes (green). Bottom panel: number of utilized threads (red) and CPU load (green).

### Issues

One issue prevented us to successfully test the version 01.14.11.27 (and earlier versions) in a cluster environment: Isaac does not respect the boundaries placed on the usage of threads and RAM.

For example, when specified to use 48 threads for reference indexing on the “high-memory” machine, Isaac still uses all available 96 threads in certain phases of computation, according to its ouput log. Similarly, Isaac aligner does not tend to respect the user-set number of threads specified to it on the command line using option -j. For example, when we ran tests on the “high-memory node”, which has 48 dual-threaded cores, with -j 48, the aligner logs still indicated that in fact all 96 virtual threads were used.

For this reason, when running on a cluster node, especially in a shared mode with other users, Isaac throws a “`libgomp: Thread creation failed”` error (found both on Biocluster and iForge). For example the Biocluster nodes in question have 384 CPUs per node, and 2 TB of RAM per node. Thus, we are not exceeding the number of cores on the node by asking Isaac aligner to use 48 threads via the –j option, and also telling the PBS script to submit the Isaac job with a limitation of 48 threads, so that other users could utilize the other threads. However, Isaac appears to ignore these limitations: not only ignoring the -j option, but also not complying with the scheduler’s limits, and throwing the “`libgomp: Thread creation failed”` error.

Isaac was probably designed to run alone on a computer, and that may be why we are seeing this error. It was reported to the developers, who did come up with a fix in January 2015. We did not test it on a cluster again.

### Summary and conclusions

In summary, the Isaac workflow appears to be optimized for whole genome sequencing on NA12878. It reports high number of false positive variants on exome data, both synthetic and real. However, its accuracy has improved steadily over several versions. On synthetic WGS, the accuracy of Isaac v2 is comparable or slightly worse than BWA-GATK and Novoalign-GATK. Special attention to variant filters is required when evaluating results.

In terms of performance, it is best to dedicate a single server to Isaac, where it will not compete with other software for hardware resources, otherwise the user might notice unstable behavior.

## Acknowledgements

We are grateful to Dr. Volodymyr Kindratenko (ISL2.0, NCSA) for allowing us to use his high memory node for the testing of Isaac workflow. Many thanks also to the system administrators at the Private Sector Program (NCSA) for their help in installing and determining reasons for runtime failures on iForge. Finally, we would like to acknowledge the Computer Network Resource Group at the Institute for Genomic Biology for similar efforts on Biocluster.

This research is part of the Blue Waters sustained-petascale computing project, which is supported by the National Science Foundation (awards OCI-0725070 and ACI-1238993) and the state of Illinois. Blue Waters is a joint effort of the University of Illinois at Urbana-Champaign and its National Center for Supercomputing Applications.

